# Quality control analysis of the 1000 Genomes Project Omni2.5 genotypes

**DOI:** 10.1101/078600

**Authors:** Nicole M. Roslin, Li Weili, Andrew D. Paterson, Lisa J. Strug

## Abstract

**Citation:** For any use of the 1000 Genomes Project data, please use the citation as noted here: http://www.1000genomes.org/faq/how-do-i-cite-1000-genomes-project. To cite this report or the lists described here, please use the following:

Roslin NM, Li W, Paterson AD, Strug LJ. Quality control analysis of the 1000 Genomes Project Omni2.5 genotypes (Abstract/Program #576/F). Presented at the 66^th^ Annual Meeting of The American Society of Human Genetics, October 18-22, 2016, Vancouver, Canada.

**Data Summary:** *Chips*: IlluminaHumanOmni2.5-4v1_B and Illumina HumanOmni25M-8v1-1_B

*Initial number of SNPs*: 2 458 861

*Initial number of samples*: 2318

*Number of SNPs passing QC*: 1 989 184 (80.9%)

*Number of samples passing QC*: 2318 (100%)

*Number of quasi-unrelated samples with consistent ethnicity and well inferred sex*: 1736

**Abstract:** The 1000 Genomes Project genotype 2318 individuals (48.1% male) from 19 populations in 5 continental groups on the Illumina Omni2.5 platform. The data are publicly available, and will prove a valuable resource to obtain ethnic-specific allele frequencies, as well as exploring population histories through principal components analysis (PCA), estimation of inbreeding coefficients, and admixture analysis. As in any study, the data should be cleaned prior to analysis, to remove individuals or markers of questionable quality. Furthermore, a thorough understanding of the relationships between individuals must be established. Here we report our findings after comprehensive examination of the data for quality control.

The basic quality of the genotypes was assessed using standard procedures. KING version 1.4 was used to confirm the relationships in the provided pedigrees, and also to detect undeclared relationships. PCA was used to examine the similarities and differences between individuals among and between population groups.

In general, the data was found to be of high quality. No samples were removed due to low call rate (<97%) or excess heterozygosity. Sex chromosome genotypes showed two individuals with discrepancies between reported and inferred sex, and were unable to determine sex in an additional 20 individuals; the sex for these was changed to unknown. Relationship checking found discrepancies between first-degree relationships in the provided pedigrees and the genotypes in 9 families, including one instance where a reported parent/child pair was unrelated, two instances where full sibs were unrelated, and one set of three individuals who formed a newly defined trio. A set of 1756 individuals who were inferred to be more distant than 3^rd^ degree relatives was extracted and used in PCA. These individuals clustered in a pattern that is consistent with other published reports of global populations. We identified 4 individuals whose genotypes clustered more closely with a different geographic region than the one in the provided data.

Although the genotype data is of high quality, errors exist in the publicly available dataset that require attention prior to using the genotypes. PLINK-format files including SNPs with good quality metrics and revised pedigree structures is available at http://tcag.ca. Files with distantly related or unrelated individuals, with sex inference consistent with provided gender, and with PCA consistent with continental group are also available.

## 1 Background

Genotypes generated on the Illumina Omni2.5 platform are available for download from the 1000 Genomes Project website (http://www.1000genomes.org/). This data is a valuable resource of genotypes from individuals collected from multiple ethnic groups around the world without regard to disease. The data can be used either as representative genotypes from individuals of known ethnicity for population structure analysis, or as a source of ethnic-specific allele frequencies which can be used in analyses such as linkage analysis, relationship estimation and estimation of inbreeding coefficients. As in any study, the data should be cleaned prior to analysis, to remove individuals or markers of questionable quality.

Furthermore, a thorough understanding of the relationships between individuals must be established. This report describes the quality analyses that we performed on the chip data. Lists of SNPs and samples with good quality metrics are available at http://tcag.ca/. This analysis was approved by the Research Ethics Board at The Hospital for Sick Children (REB number 1000054008).

## 2 Data acquisition

Genotype files, in vcf format, were downloaded on 16 May 2016 from the following ftp site at the 1000 Genomes Project: ftp://ftp.1000genomes.ebi.ac.uk/vol1/ftp/release/20130502/supporting/hd_genotype_chip/ (Genomes Project et al., 2015). The link to this directory came from the Frequently Asked Questions section of the website (http://www.1000genomes.org/faq/what-omni-genotype-data-do-you-have/). Genotypes were generated at the Broad and Sanger Institutes, using two different versions of the Omni2.5 platform. The majority of the samples were genotyped at the Broad, and no individuals were genotyped in both locations. The analysis presented in this report started with the downloaded file `ALL.chip.omni_broad_sanger_combined.20140818.snps.genotypes.vcf.gz`, which is the combined set of Broad and Sanger genotypes. The data had 2318 individuals and 2 458 861 markers. See Appendix 2 for the specific details regarding the downloaded files.

## 3 Prior processing by 1000 Genomes

Before the data was made available for download, the data providers performed a small amount of processing on the data, which is summarized here. For the data generated at the Broad, genotypes were called using GenomeStudio v2010.3 with the calling algorithm/genotyping module version 1.8.4 using the default cluster file. Physical positions were taken from build 36, which were subsequently converted to hg19 using the human_g1k_v37 dataset. SNPs were removed according to the following filters: the metadata on the chip for the SNP was inconsistent, the SNP had a duplicate with a higher Gentrain Score, the assay was designed for multiple alternate alleles, the SNP was not polymorphic in the 1000 Genomes dataset, or the SNP was within 50bp of a known indel. These filters resulted in 2 441 898 SNPs remaining (out of 2 450 000, 99.7%). The specific chip used was IlluminaHumanOmni2.5-4v1_B.

For the data generated at the Sanger Center, detailed information on prior processing was not provided. However, of the 2 391 739 SNPs on the chip, genotypes were provided for 2 176 330 (91.0%). Note that rs numbers were not provided for the variants in this dataset. It was assumed that the physical positions were based on hg19. The genotypes came from the IlluminaHumanOmni25M-8v1-1_B chip.

The two datasets were merged using GATK v3.1.1 combineVariants. The final dataset had 2318 individuals and 2 458 861 markers. No individuals were present in both sets of data. The majority of SNPs (2 150 028, 87.4%) were present in both the Broad and Sanger datasets; 282 531 (11.5%) SNPs were present just in the data generated at the Broad, while 26 302 (1.1%) SNPs were only in the data from the Sanger. In the final dataset, there were no markers with identical chromosome and physical positions. The merged data was saved to file Omni25_sanger_broad_combined.vcf.gz and made available on the 1000 Genomes ftp site.

## 4 Family and ethnic information

Gender and basic information on relationships between certain individuals was provided by 1000 Genomes in a pedigree file (Appendix 2, file 2). Additionally, reported ethnic background was provided as a file from each genotyping centre (Appendix 2, files 3 and 4). The pedigree file included only individuals genotyped or sequenced in the 1000 Genomes project, and so additional "dummy" individuals needed to link up related individuals were not present. Because of this, several known relationships were not made explicit by the pedigree. All samples from Sanger and 2098 individuals from the Broad had available ethnic information, while 43 individuals genotyped at the Broad had missing ethnicity and gender in the provided files, for a total of 2318 individuals. Missing genders and ethnicities were filled in using file 5 in Appendix 2. Counts of individuals from each of the sampled populations are shown in Appendix 1.

## 5 Quality control (QC) analysis

The provided vcf files were converted to PLINK binary format using PLINK1.90 (https://www.cog-genomics.org/plink2/). One marker had a non-valid chromosome code (chromosome GL000202.1 for SNP11-69436716); the chromosome for this marker was set to missing. The provided sex and relationship information were incorporated into the pedigree files. These files were used as the basis for the QC steps that follow. Analyses were performed using in-house scripts, unless otherwise stated.

### 5.1 Basic sample quality

Analyses were performed to determine the general quality of the samples. Since the two different genotyping centres used slightly different sets of markers, only the 2 150 028 SNPs present in both the Broad and Sanger datasets were used in these tests.

#### 5.1.1 Sex chromosome analysis

Sex chromosome composition was inferred for each of the samples based on the heterozygosity rate on chromosome X and the call rate on chromosome Y. The majority of samples could be clearly identified as XY or XX, according to the following criteria:

1. XY (male) = proportion of heterozygous genotypes on the X chromosome >0.9, and call rate on the Y chromosome >0.9.
2. XX (female) = proportion of heterozygous genotypes on the X chromosome <0.9, and call rate on the Y chromosome <0.4.

All other individuals were declared to have ambiguous (or unknown) sex. Based on these thresholds, two individuals, NA21310 and HG02300, were listed as males, but had genotypes consistent with females (Figure 1). No other sex discrepancy was found; however, for 20 individuals, the sex could not be inferred using the above rules. The sex for these individuals was set to missing. For 8 of these individuals (reported to be male), the heterozygosity rate on the X chromosome was consistent with males, but showed reduced call rate on the Y chromosome. Such a loss of Y could be the result of cell line artifacts or environmental factors (Dumanski et al., 2015), rather than genotyping or sample error, and so these individuals were not flagged as having poor quality. Similarly, 11 reported females showed excessive homozygosity on chromosome X, also consistent with cell line artifacts. One individual (HG01683: male, IBS) had slightly reduced call rate on Y for a male, but X chromosome heterozygosity similar to females, suggesting the possibility of XXY.

**Figure 1.**
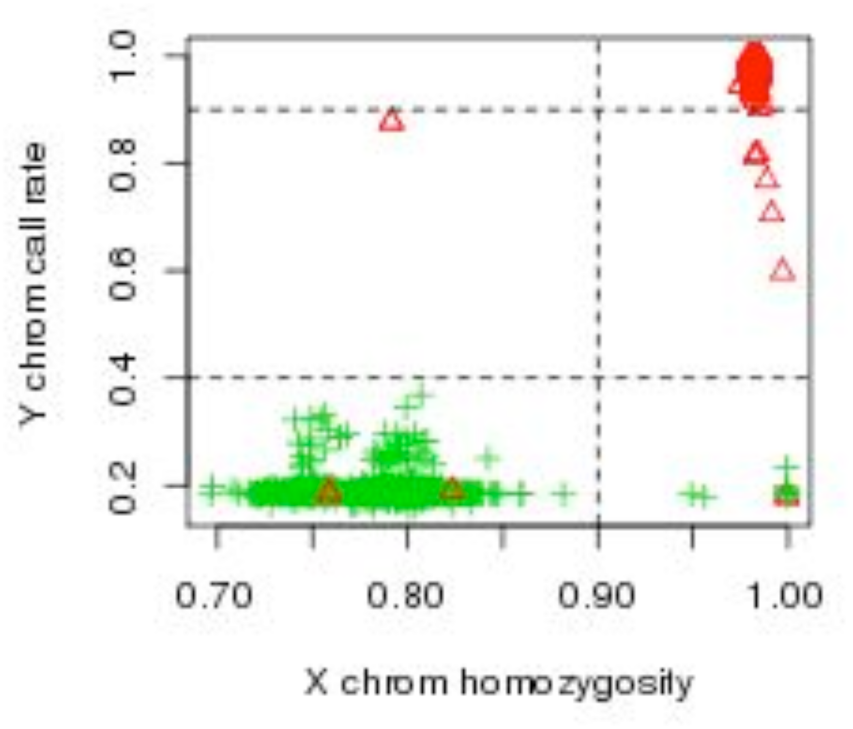
Proportion of homozygous genotypes on the X chromosome vs. proportion of non-missing genotypes on the Y chromosome. Symbols are coded based on the sex in the provided pedigree file: male = red triangles, female = green plus symbols. Dashed lines delimit the thresholds applied above.

#### 5.1.2 Call rate

The proportion of non-missing genotypes (call rate) was calculated per individual. In general, the call rate was very high, with an average call rate of 99.6% in the data. The lowest call rate per sample was 97.0%, which is at the commonly-accepted threshold of 97%. Call rates were calculated using PLINK1.90.

**Figure 2.**
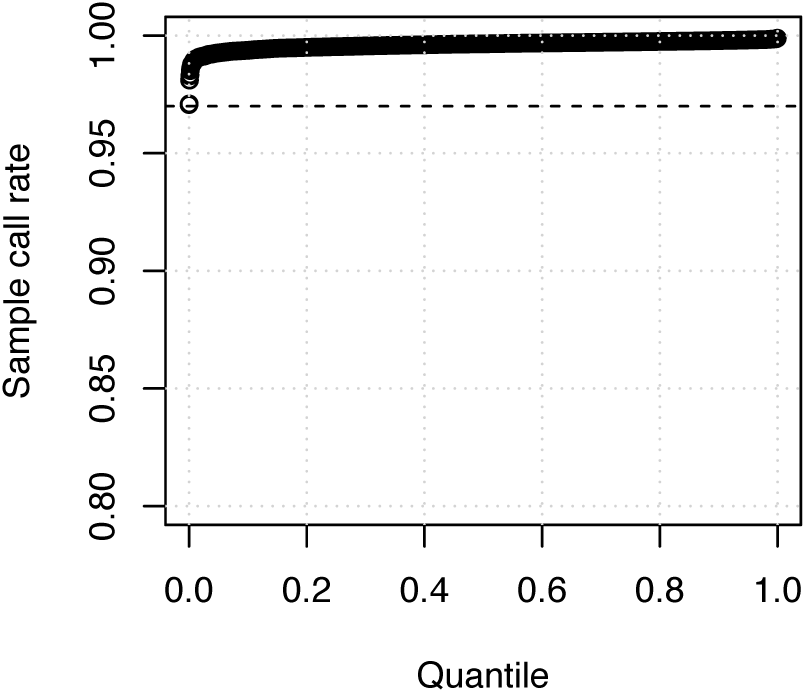
Proportion of non-missing genotypes per individual. The dashed line shows the 97% threshold.

#### 5.1.3 Multilocus heterozygosity

In order to detect possible sample contamination, the average autosomal heterozygosity was calculated for each individual. The average heterozygosity was 0.20, and no sample was unusually extreme, compared to the other samples (Figure 3). The wide distribution with multiple peaks are likely reflections of the multi-ethnic nature of the genotyped individuals.

**Figure 3.**
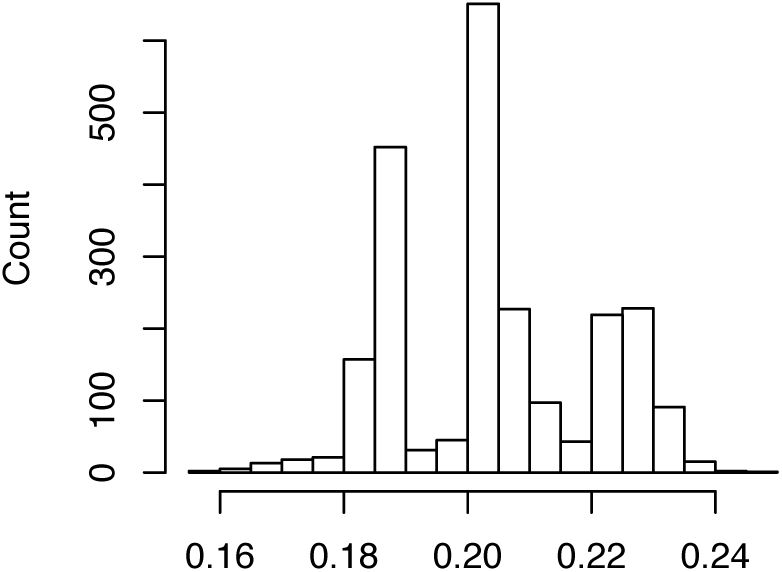
Histogram of the proportion of heterozygous autosomal loci per sample.

#### 5.1.4 Heterozygous haploid genotypes

Haploid regions of the genome are typically represented as homozygous diploid genotypes. These regions include the X chromosome for males, the Y chromosome, and the mitochondrion. Individuals with an unusually high number of heterozygous genotypes at these loci could provide additional evidence of quality issues. Since the regions to examine depend on sex, the sex inferred above was used, rather than the provided sex. In the current dataset, no individual had an unusually high number of heterozygous haploid genotypes, relative to the other samples (results not shown). These genotypes were detected using PLINK.

#### 5.1.5 Summary of basic sample quality

In general, the quality of the samples was excellent. There was no evidence of contamination, and no evidence of sample failure. Inferred sex was consistent with provided sex for 2296 of 2318 (99%) of the individuals. The sex for two individuals (NA21310 and HG02300) was changed from male to female, while the sex for 20 individuals was set to missing.

### 5.2 Basic quality per SNP

Standard tests were performed to determine SNP quality. Only markers that were present on both versions of the chip were included–the other markers will have non-random sets of missing values, which will cause bias in many of the tests. The analyses were performed after the sex changes described above.

#### 5.2.1 Call rate

The proportion of non-missing genotypes was calculated per SNP. The call rate was excellent: 2 114 511 of 2 150 028 (98.3%) of the SNPs had call rate >97% (Figure 4).

**Figure 4.**
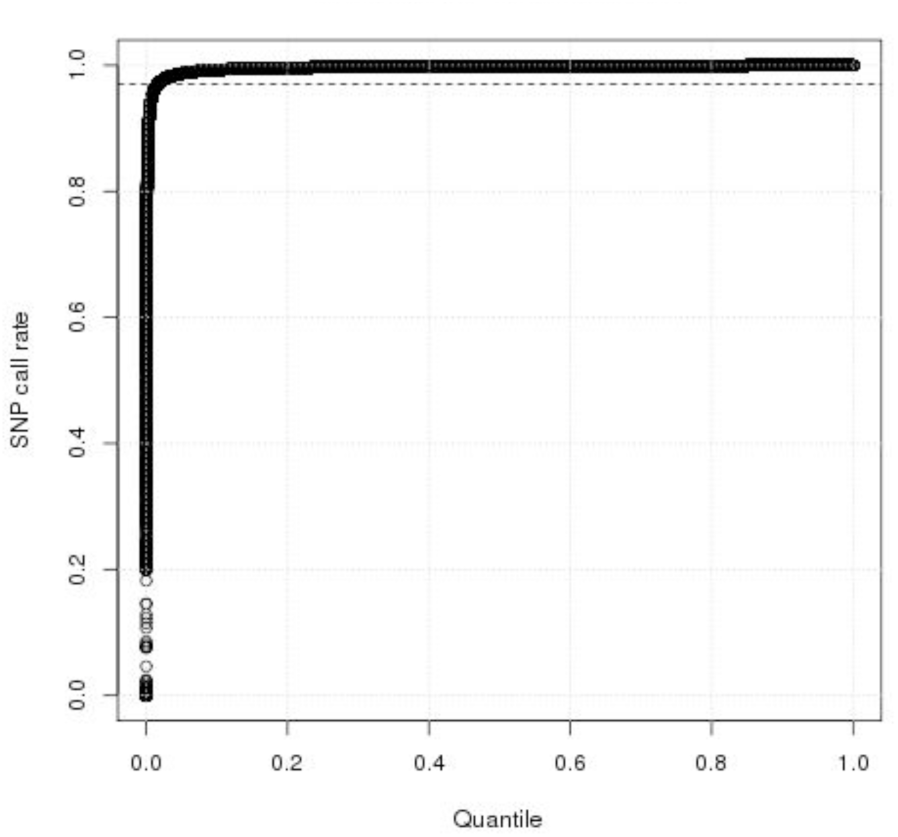
Call rate per SNP, for the markers present in both the Broad and Sanger datasets. The dashed line shows the 97% threshold.

#### 5.2.2 Heterozygous haploid (HH) genotypes

The distribution of heterozygous haploid genotypes, defined in section 5.1.4 above, was examined. Non-missing genotypes for females on the Y chromosome were also identified. A total of 11 888 SNPs had at least one HH genotype. No SNP appeared to have an excessive number of these problematic genotypes, compared to the rest of the SNPs (results not shown).

#### 5.2.3 Hardy-Weinberg equilibrium

SNPs were tested for deviation from the proportions expected under Hardy-Weinberg equilibrium (HWE), using an exact test (Wigginton, Cutler, & Abecasis, 2005) implemented in PLINK. For SNPs on the X chromosome, only inferred females were used in the test. SNPs on the Y chromosome and the mitochondrion were not tested. A total of 355 357 SNPs (17%) had p-values <10^−6^. Although this could indicate substantial problems in the allele calling algorithm, it is more likely due to the fact that the samples came from a wide variety of ethnic backgrounds, and the assumptions of Hardy-Weinberg do not hold (most importantly the assumption of random mating) in the data. Therefore, SNPs were not excluded due to deviation from HWE.

#### 5.2.4 Summary of basic SNP quality

The SNPs had very high quality. SNPs were removed if the call rate was <97% or if a HH genotype was detected. Furthermore, SNPs that were monomorphic in all the samples were discarded, since they do not contain any information. Out of the original 2 150 028 markers that were present in both the Broad and Sanger datasets, 2 105 791 (97.9%) passed these quality filters, and will be used in the more sophisticated analyses below.

### 5.3 Relationship testing

The software KING 1.4 (Manichaikul et al., 2010) was used to infer close relationships between pairs of samples. This program estimates kinship coefficients for all pairs of individuals based on their heterozygosity, and does not require allele frequency estimates. The high quality set of 2 105 791 SNPs described above was used. KING was also used to select a set of individuals who were inferred to be unrelated or more distantly related than 3^rd^ degree (first cousin or equivalent).

Prior to testing, the relationship information provided by 1000 Genomes was incorporated. There were 28 pedigrees that were larger than trios, shown in Appendix 3, plus an additional 373 trios (based on the number of individuals per family ID). In many cases, larger families were composed of multiple smaller families with different pedigree IDs. We assigned these a new family ID based on the population from which the families came. For example, the three individuals HG00501 (a mother in a trio, family SH028), HG00512 (a father in a trio, family SH032) and HG00524 (a father in a trio, family SH036) were all listed as siblings, even though they had different family IDs. We combined all thee trios into a single family, adding dummy parents, and named the extended family CHS3.

Within the 28 pedigrees, the pairwise kinship coefficients were consistent with the provided structure for 23 of them. For the remaining 5 families, ASW3 was split into two separate families (there may have been a typo in the provided pedigree information, mixing up IDs NA20334 and NA20344). Families ASW4 and ASW5 were combined and one half sibling changed fathers. Family LWK003 originally consisted of two individuals with a common mother, with no information on the father(s). The genotypes were consistent with a full sibling relationship, and so a single dummy father was created. Finally, family YRI4 was split into two unrelated families.

In the trio CLM23, the kinship coefficient between the parents was estimated to be 0.033, indicating that they may be distantly related (consistent with first cousin once removed, or equivalent relationship). Since there are no other family members with genotypes available, it is not possible to confirm this relationship, and so the pedigree was not changed. A paper estimated inbreeding coefficients in 1000 Genomes samples based on whole genome sequence data (Gazal, Sahbatou, Babron, Genin, & Leutenegger, 2015), including the parents of trio CLM23. The father of the trio (HG01277) was found to have an inbreeding coefficient estimate consistent with offspring of second cousins. However, the child in the trio (HG01279) was not analyzed, and so this analysis cannot confirm a relationship between the parents.

Other than the changes made to the extended families, 5 additional changes were made, based on inferred first-degree relationships between pairs of individuals with different family IDs. There were 3 instances of unreported full sibling relationships, one instance where three supposedly unrelated individuals formed a trio, and one instance where a reported parent/offspring relationship was inferred to be unrelated. A substantial number of 2^nd^ degree relationships were also observed, but pedigrees were not modified based on these more distant kinship coefficients. All pedigree changes are shown in Appendix 4.

KING was used to select quasi-unrelated individuals. It identified 1756 individuals who were no closer than 3^rd^ degree relatives (first cousin or equivalent).

#### 5.3.1 Mendelian errors

The program PEDSTATS 0.6.10 (Wigginton & Abecasis, 2005) was used to look for errors in Mendelian transmission in the revised pedigrees identified above. Only the autosomes and chromosome X were examined. A total of 228 542 errors were found. Out of the pedigrees having at least one parent/child pair, family 1349 had the most errors (5743). Since over 2 million SNPs were tested, this represents errors in approximately 0.2% of the markers, and so does not present a quality concern. The distribution of errors by SNP was also not remarkable (results not shown). All 116 607 SNPs with ≥1 error were removed, resulting in 1 989 184 SNPs.

### 5.4 Principal components analysis

Principal components analysis (PCA) was used as a visual aid to show how similar the genotypes from the data were, when the provided ethnic information was included. In order to ease computational burden, and to avoid artifacts from unusual patterns of linkage disequilibrium (LD) that are unrelated to ethnicity, the set of 1 989 184 SNPs was further reduced. First, all markers on chromosome X were removed. Next, two regions with unusual patterns of linkage disequilibrium (LD) were removed: the major histocompatibility region on chromosome 6, and an inversion polymorphism on chromosome 8. Positions on the genetic map were obtained from the Rutgers combined map (Matise et al., 2007), and only markers with unique cM positions (up to 3 decimal places) were retained. Arbitrarily, out of a group of SNPs at the same genetic position, the one with the lowest physical position was retained. Markers with minor allele frequency (MAF) >0.3 in the complete set of data were extracted. This resulted in a set of 282 273 markers. To reduce tight LD between markers, the set of SNPs was further reduced so that all markers had pairwise r^2^ <0.2 within each chromosome, to produce a final set of 57 931 SNPs. Since close relationships between individuals can distort the PCA results, only the 1756 quasi-unrelated individuals identified above were used in the analysis. PCA was performed on this data using the SmartPCA package (version 10210) of Eigenstrat (Patterson, Price, & Reich, 2006; Price et al., 2006).

Although only a subset of individuals recruited for the 1000 Genomes Project have Omni2.5 genotypes, individuals from all continental groups are represented. The distribution of samples in the PCA is consistent with what has been shown elsewhere, with AFR, EAS and EUR forming major axes, and AMR and SAS lying within these axes (Figure 5). Four individuals did not cluster well with other individuals from their continental groups. Three individuals from AMR (HG01241, HG01242 and HG01108, all from PUR) clustered more closely with individuals from AFR, and one individual from AFR (NA20314, ASW) was more similar to AMR. These results are not surprising, given the population histories. The two individuals with sex discrepancy clustered well within their listed geographic region.

**Figure 5.**
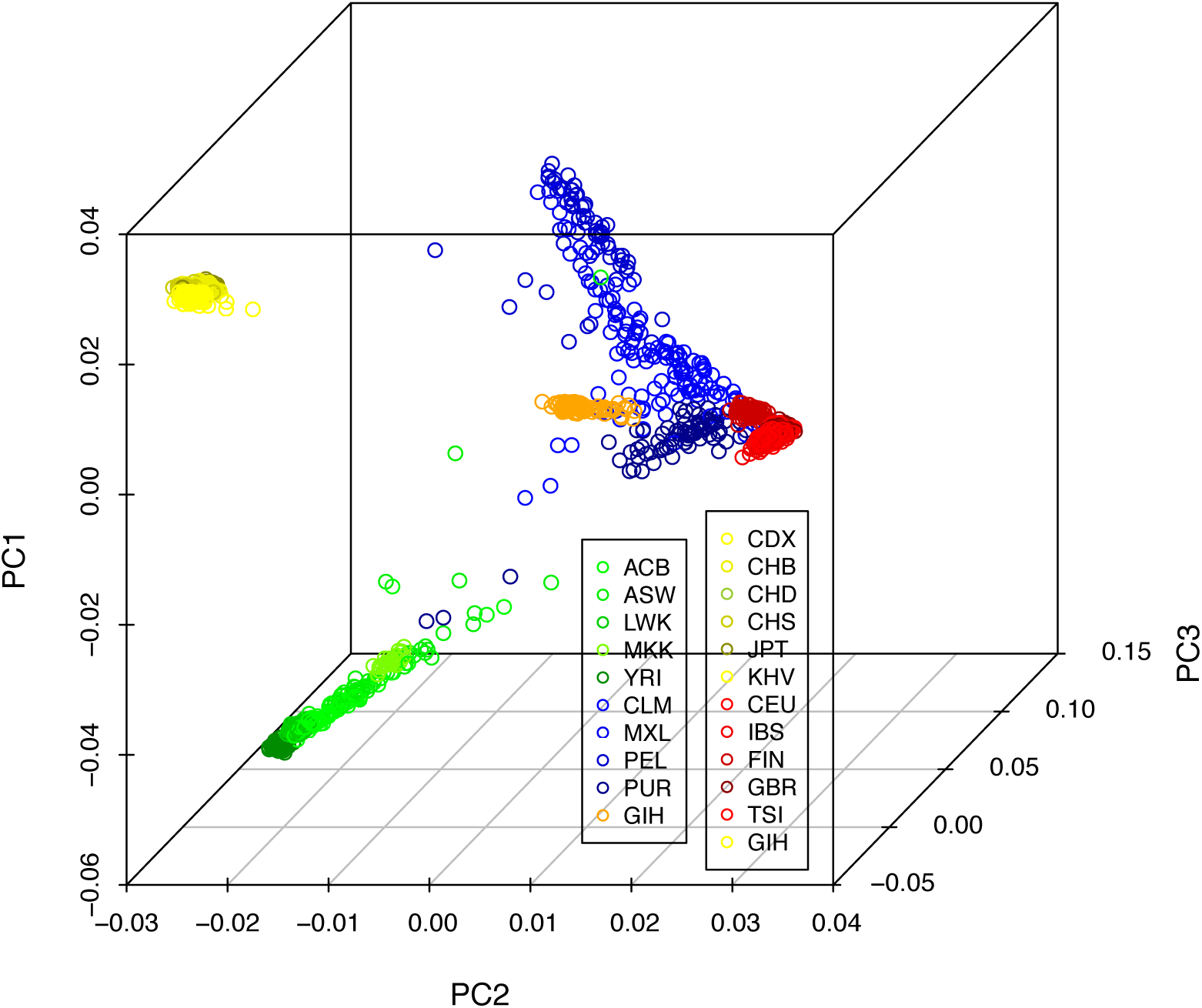
Three dimensional plot of the first three principal components (PC1, PC2, PC3) of 1756 quasi-unrelated individuals. Points are shaded by their population codes. In general, green = AFR, blue = AMR, yellow = EAS, red = EUR, and orange = SAS.

The 4 individuals who did not cluster well with their geographic region were removed, and clustering was repeated within each geographic region. For AFR, LWK, MKK and YRI each formed tight clusters, while ACB and ASW were more spread out (Figure 6).

**Figure 6.**
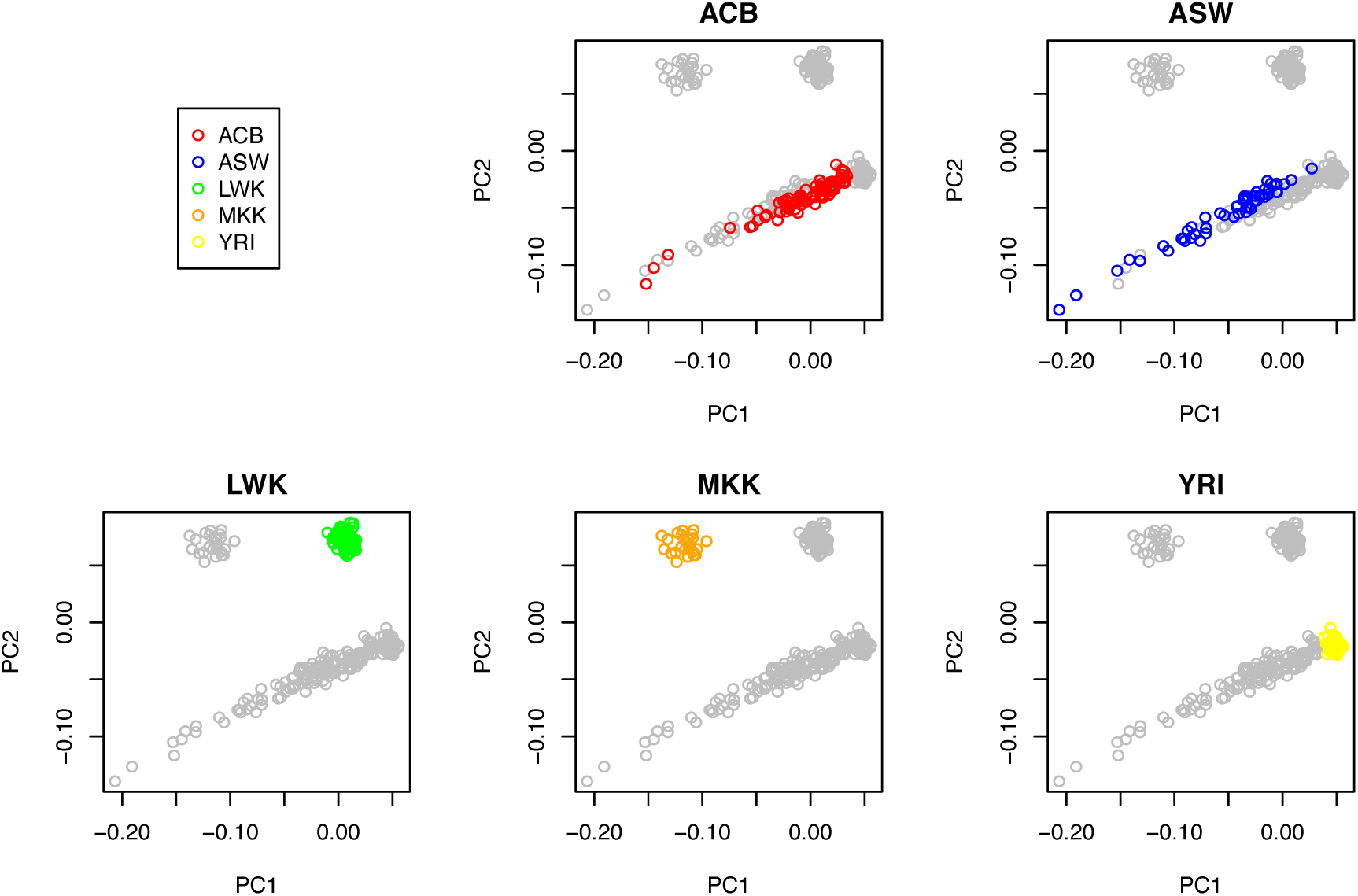
Plots of the first two PCs from an analysis of AFR. Each plot highlights one population from this continental group.

In AMR, the 4 different populations did not form tight clusters (Figure 7). Also, even though they could be differentiated somewhat, they also showed substantial amounts of overlap.

**Figure 7.**
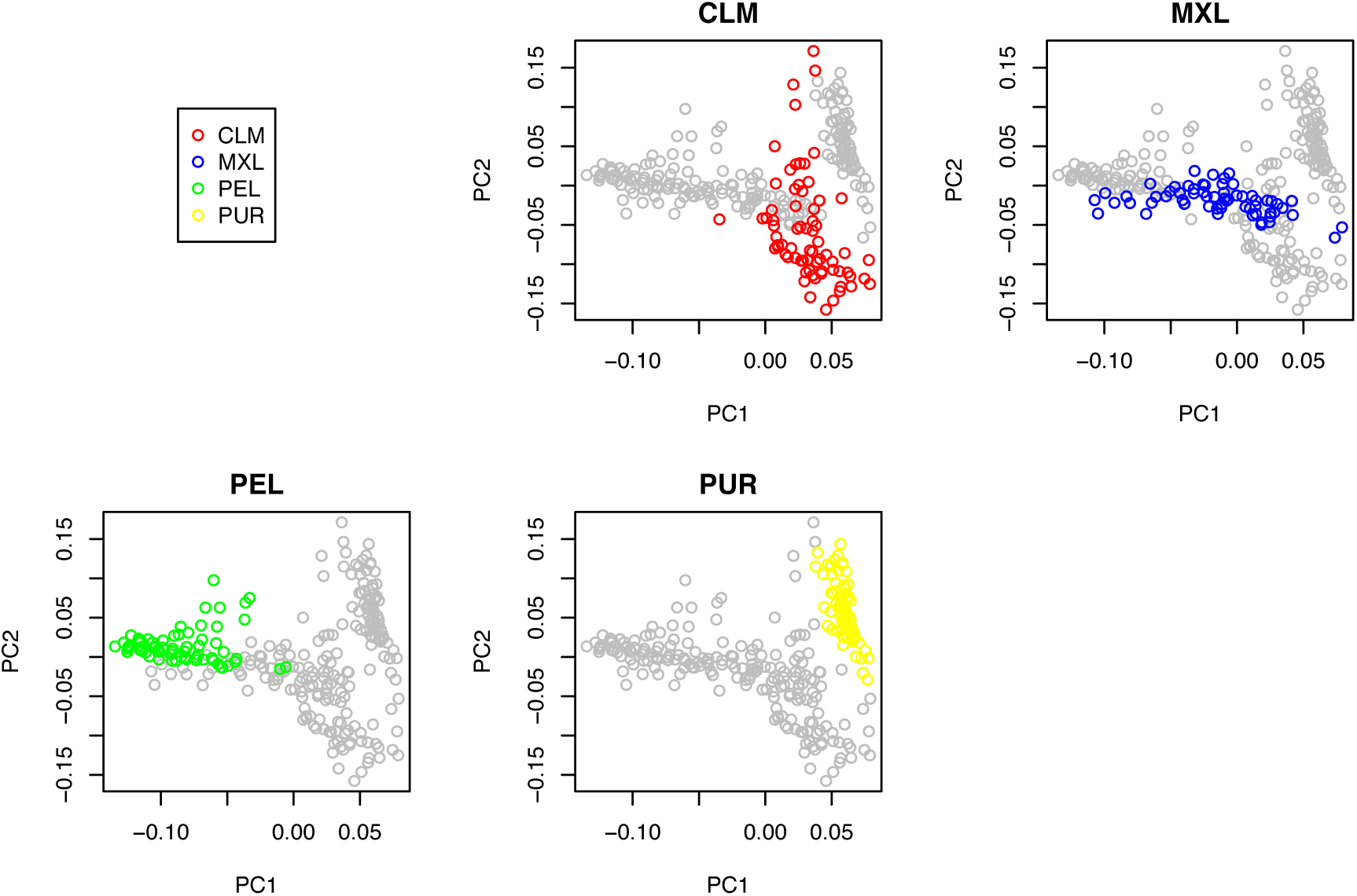
Plots of the first two PCs from an analysis of AMR. Each sub-population is highlighted in a separate plot.

Among the 5 EAS populations, JPT was distinct from CDX, CHB, CHS and KHV (Figure 8). One JPT sample may be admixed with an ancestor from a Chinese population. CDX (Chinese Dai) and KHV (Kinh from Vietnam) were more similar to each other than to the other populations from China, with the exception of one CDX individual who clustered with either CHB or CHS.

**Figure 8.**
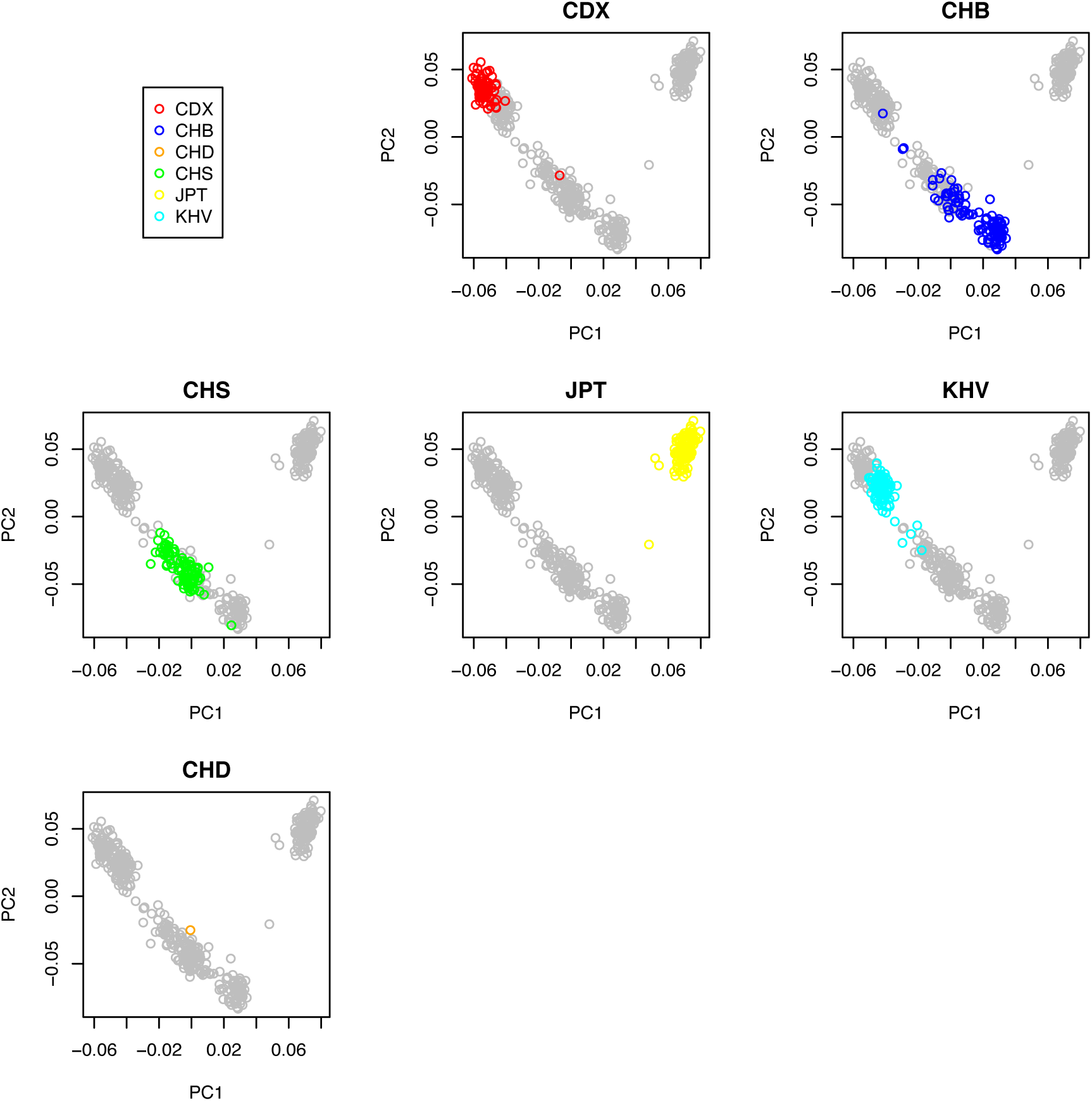
Plots of the first two PCs from an analysis of EAS. Each sub-population is highlighted in a separate plot.

The 5 populations from EUR formed three main clusters (Figure 9). CEU and GBR were indistinguishable from each other, while IBS and TSI showed similarities to each other. FIN was distinct from the other 4 populations.

**Figure 9.**
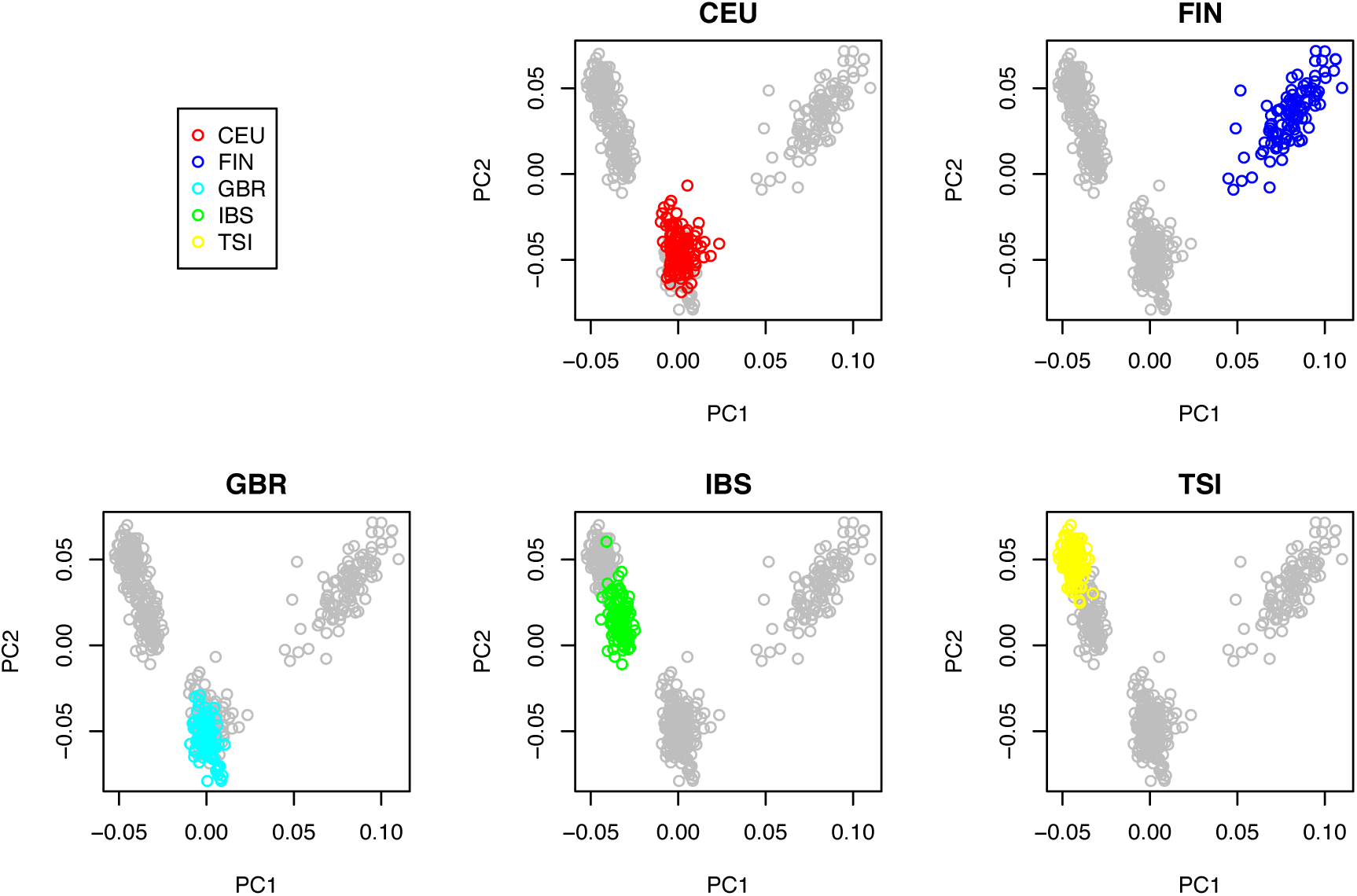
Plots of the first two PCs from an analysis of EUR. Each sub-population is highlighted in a separate plot.

Since only one ethnic group from SAS was included in the Omni2.5 genotyping (GIH), PCA was not performed within this continental group.

In total, there were 1752 individuals who were unrelated (up to 3rd degree), and who clustered well within their assigned geographic region.

## 6 Summary

The Omni2.5 genotypes provided by the 1000 Genomes Project were analyzed for quality. In general, the quality of the data was excellent. Out of 2318 samples, two individuals had genotypes on the X and Y chromosomes that were inconsistent with the provided gender, while sex could not be determined for another 20 individuals. It is recommended that these 22 individuals be removed for any analysis in which sex is important. All samples had call rate >97%. No sample had unusually high heterozygosity across the autosomes. Over 98% of the SNPs had call rate >97%.

Dummy individuals were added to the provided families to form "complete" pedigrees. Based on pairwise relationship testing, the pedigrees were modified to be consistent with first-degree estimated kinship coefficients. Changes were made to a total of 10 pedigrees.

A set of unrelated or distantly related individuals was also chosen. These individuals were more distant than 3^rd^ degree relatives, clustered with other samples from their continental group, and had clear sex inference that was consistent with the provided gender. This set had 1736 individuals.

A set of 1 989 184 high quality SNPs was selected. These markers were present in the datasets from both the Broad and Sanger Institutes, had call rate >97%, had two observed alleles, and no observed errors in Mendelian transmission.

The following files are available at our website (http://tcag.ca):

- PLINK binary format files including the 1 989 184 SNPs passing QC with two alleles, and the inferred pedigree structures, including all necessary dummy individuals
- PLINK binary format files including the 1 989 184 SNPs passing QC with two alleles, and the 1736 individuals who were inferred to be <3^rd^ degree relatives, clustered well with other individuals in their geographic region, and had clear sex inference that was consistent with provided gender
- Table of quality statistics per sample
- Table of quality statistics per SNP

## Appendix 1

*List of 1000 Genomes populations and descriptions (number of individuals with Omni2.5 data)*

<u>AFR (Africans, 508)</u>
ACB = African Caribbean in Barbados (102)
ASW = African ancestry in Southwest US (104)
LWK = Luhya in Webuye, Kenya (116)
MKK = Maasai in Kinyawa, Kenya (31)
YRI = Yoruba in Ibadan, Nigeria (189)
<u>AMR (Americas, 418)</u>
CLM = Colombian in Medellin, Colombia (107)
MXL = Mexican ancestry in Los Angeles, California (103)
PEL = Peruvian in Lima, Peru (105)
PUR = Puerto Rican in Puerto Rico (111)
<u>EAS (East Asians, 587)</u>
CDX = Chinese Dai in Xishuangbanna, China (100)
CHB = Han Chinese in Bejing, China (108)
CHD = Chinese in Denver, Colorado (1)
CHS = Southern Han Chinese, China (153)
JPT = Japanese in Tokyo, Japan (105)
KHV = Kinh in Ho Chi Minh City, Vietnam (121)
<u>EUR (Europeans, 649)</u>
CEU = Utah residents with Northern and Western European ancestry (183)
FIN = Finnish in Finland (100)
GBR = British in England and Scotland (104)
IBS = Iberian populations in Spain (150)
TSI = Toscani in Italy (112)
<u>SAS (Southern Asians, 113)</u>
GIH = Gujarati Indian in Houston, TX (113)

## Appendix 2

*List of downloaded files*

1. ftp://ftp.1000genomes.ebi.ac.uk/vol1/ftp/release/20130502/supporting/hd_genotype_chip/ALL.chip.omni_broad_sanger_combined.20140818.snps.genotypes.vcf.gz (last modified August 18 2014; merged genotype file, in vcf format)
2. ftp://ftp.1000genomes.ebi.ac.uk/vol1/ftp/technical/working/20140502_sample_summary_info/20140502_all_samples.ped (last modified May 2, 2014; sex and relationship information)
3. ftp://ftp.1000genomes.ebi.ac.uk/vol1/ftp/release/20130502/supporting/hd_genotype_chip/broad_intensities/omni25.2141.sample.panel (last modified November 22 2013; population information for samples genotyped at the Broad)
4. ftp://ftp.1000genomes.ebi.ac.uk/vol1/ftp/release/20130502/supporting/hd_genotype_chip/sanger_intensities/sanger_omni_chip.20130805.ALL.panel (last modified August 12 2013; population information for samples genotyped at Sanger)
5. http://www.1000genomes.org/sites/1000genomes.org/files/documents/20101214_1000genomes_samples.xls (population information for 43 individuals missing from Broad samples population file)

## Appendix 3

*Pedigrees larger than trios based on the information provided by 1000 Genomes; original pedigree identifiers are shown in non-black colours*

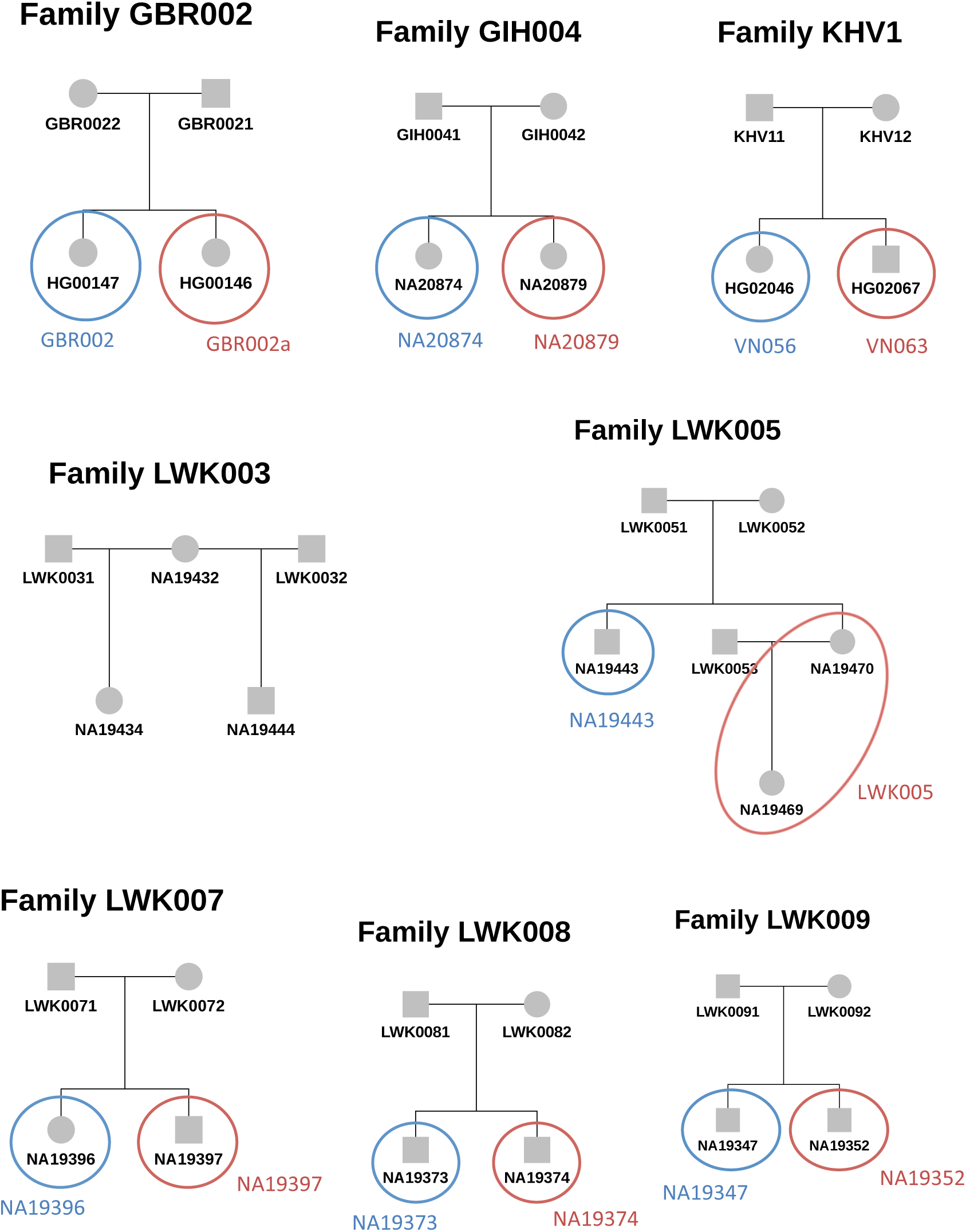

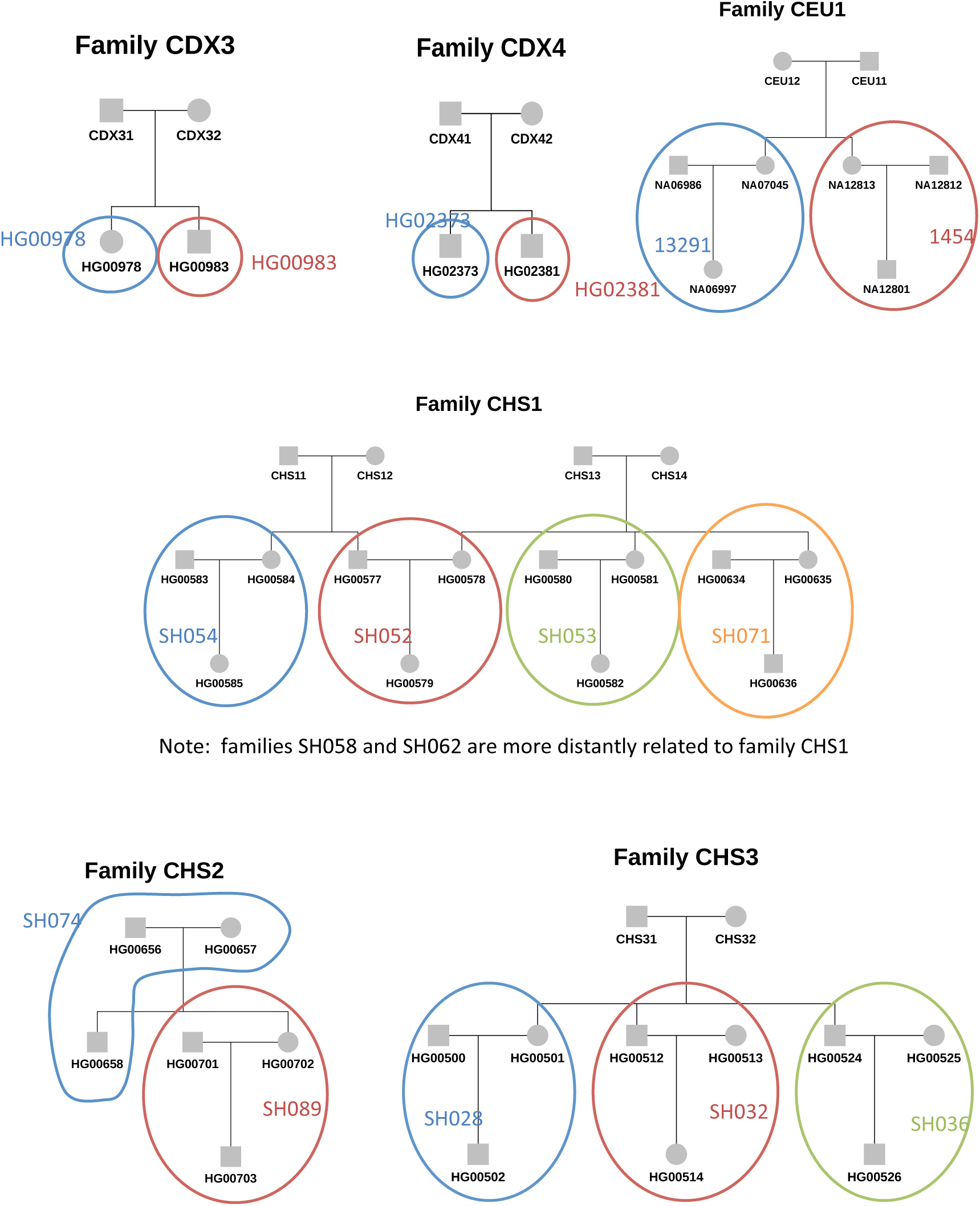

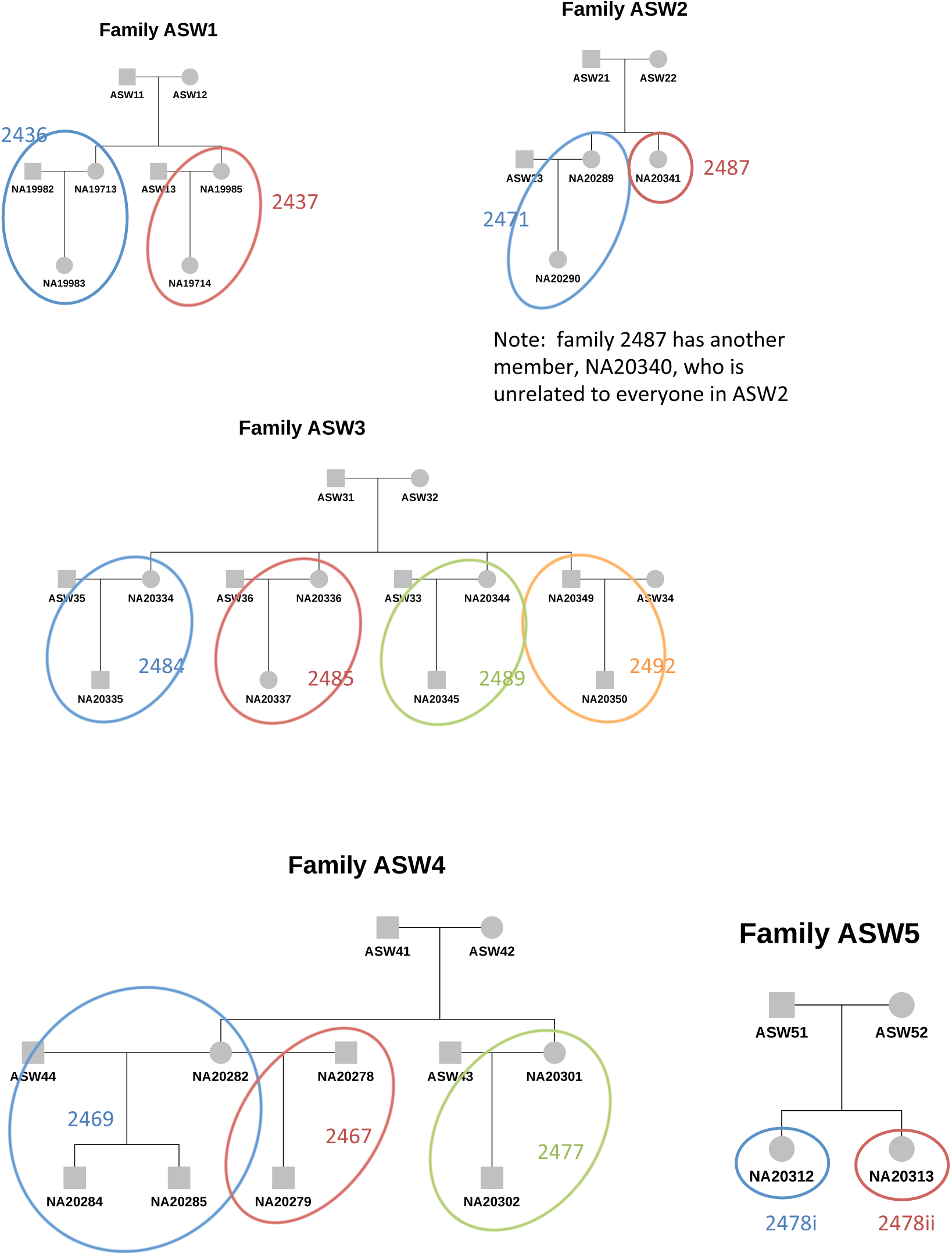

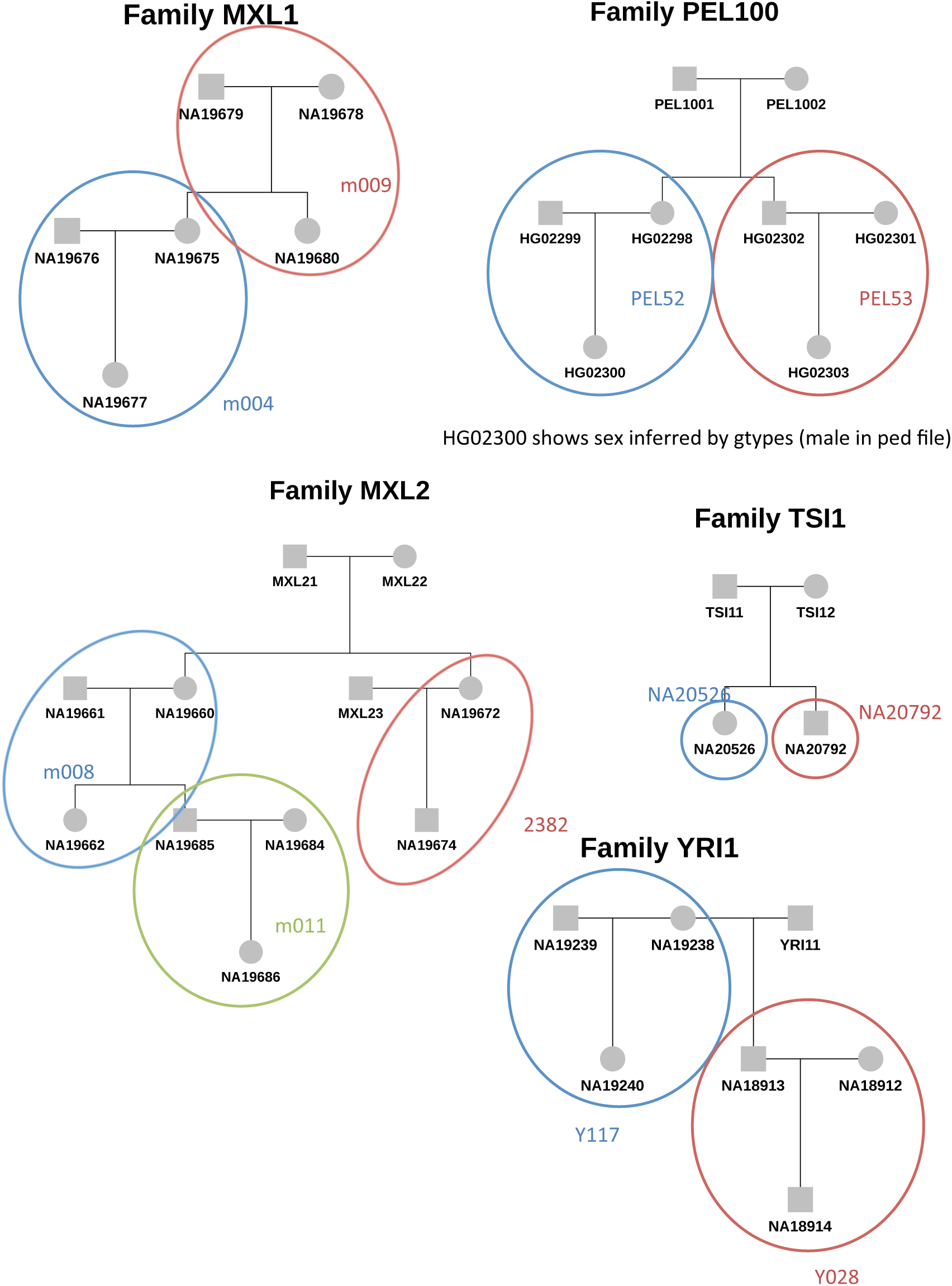

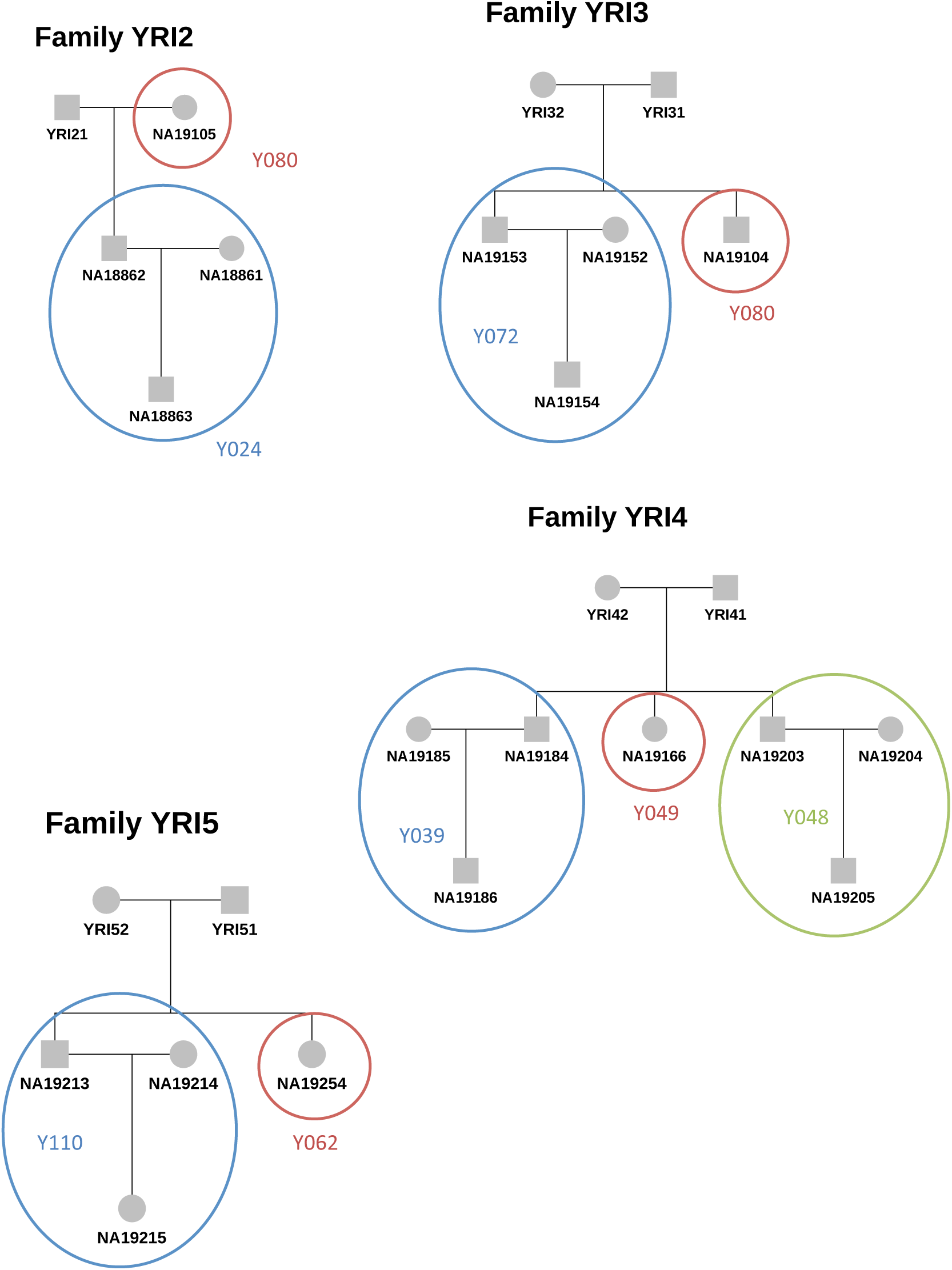

## Appendix 4

*Pedigrees that were modified after relationship inference in KING. Pedigrees prior to testing are shown in blue boxes, while pedigrees after testing are shown in red boxes.*

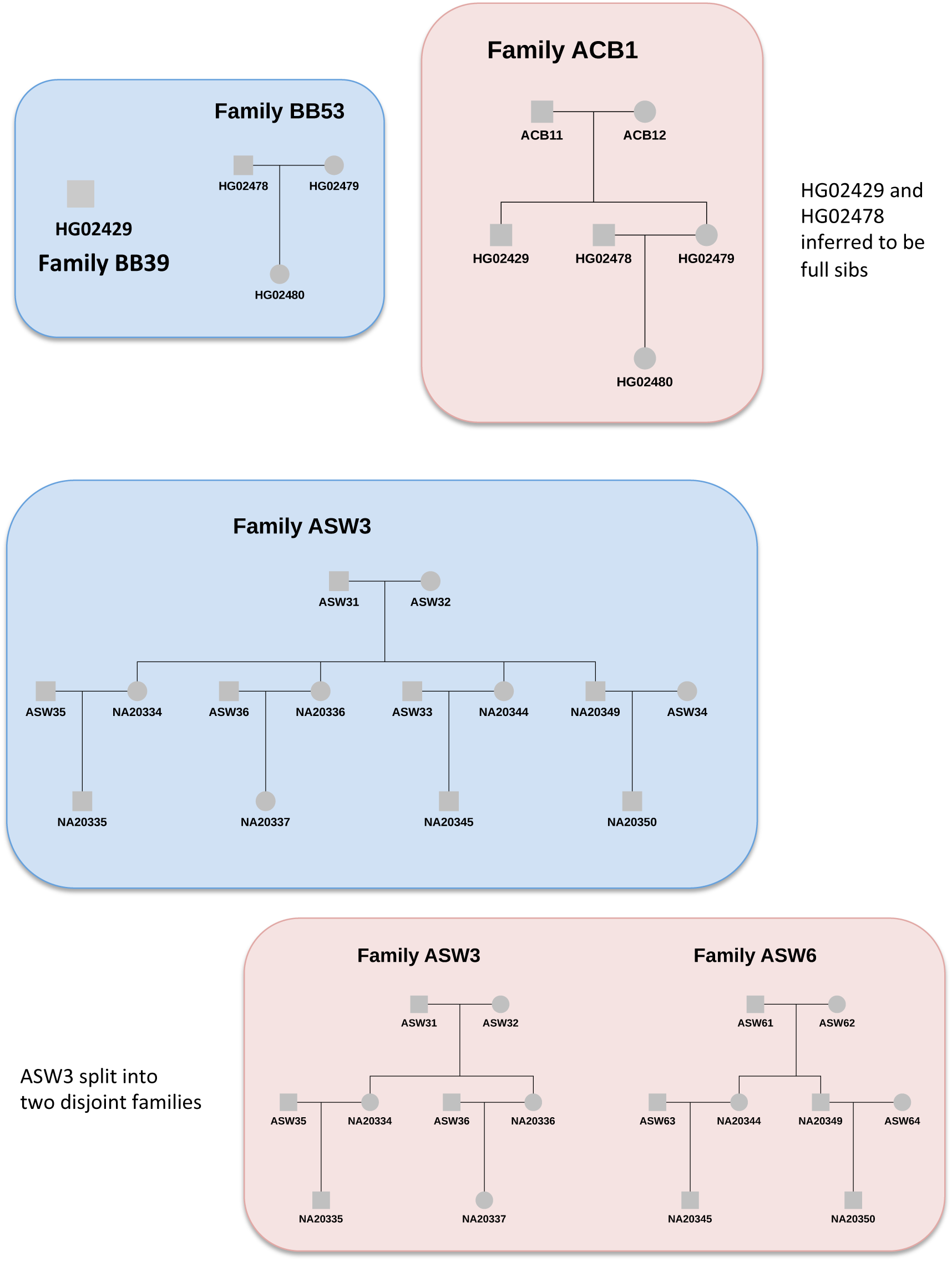

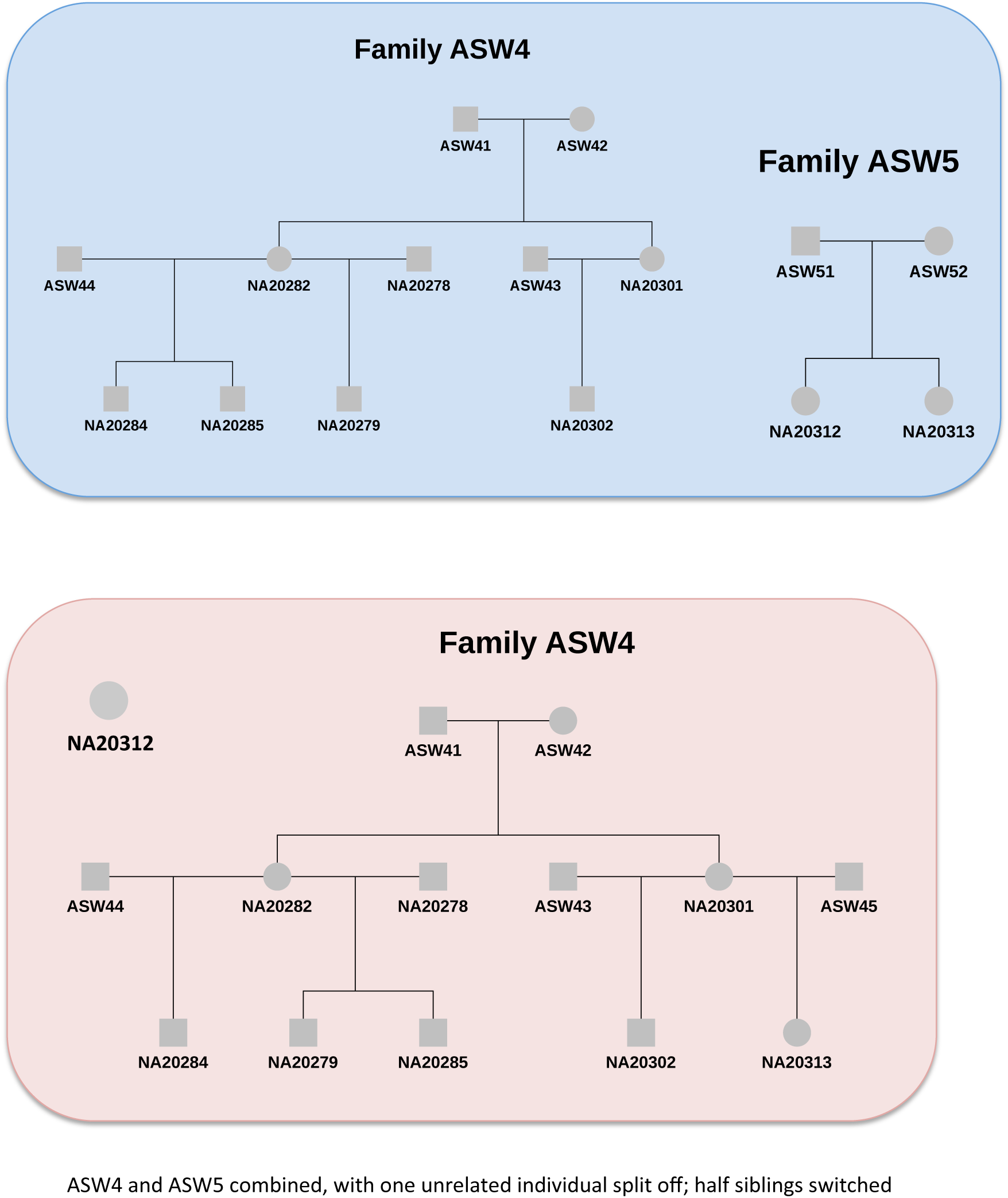

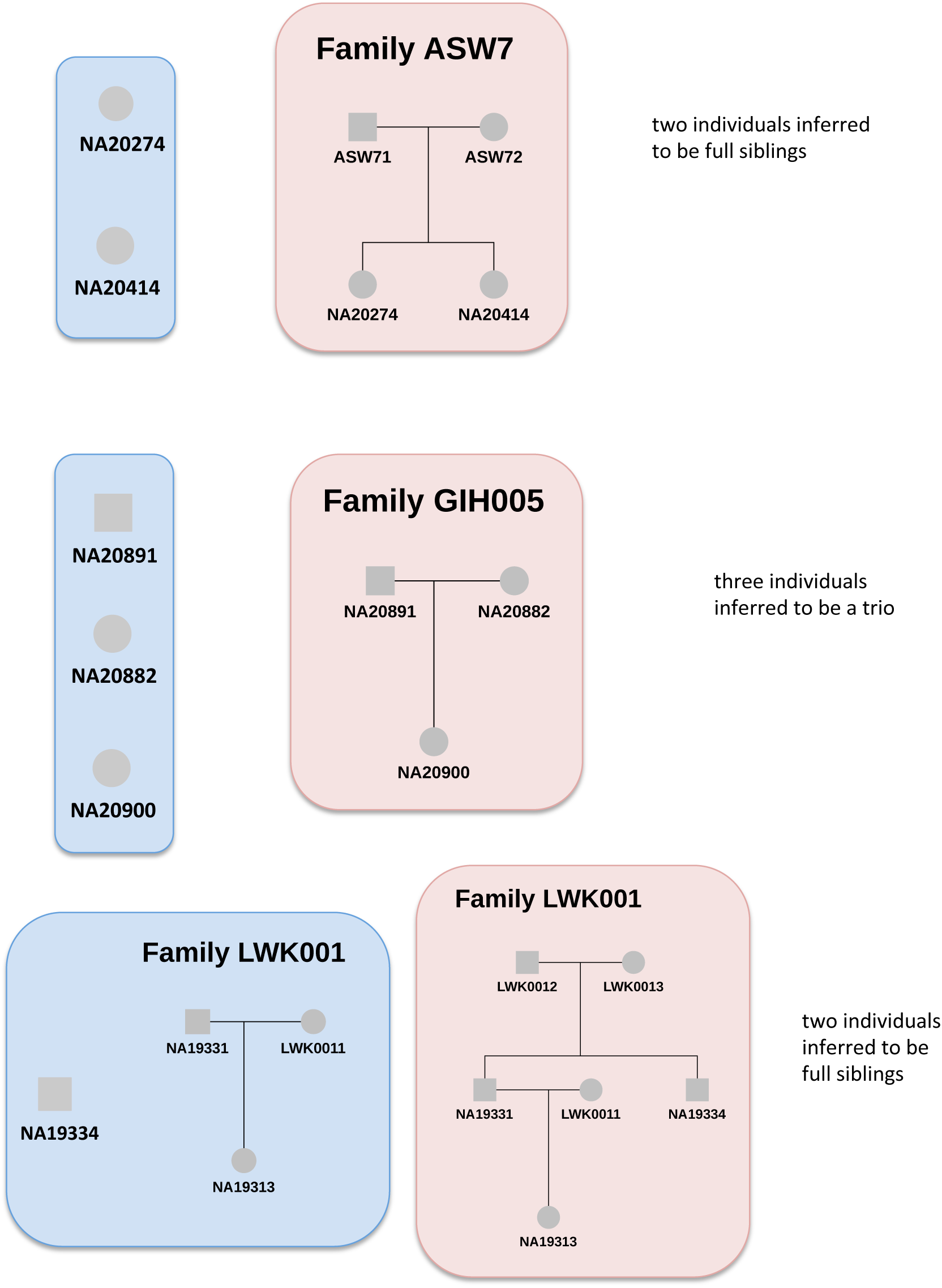

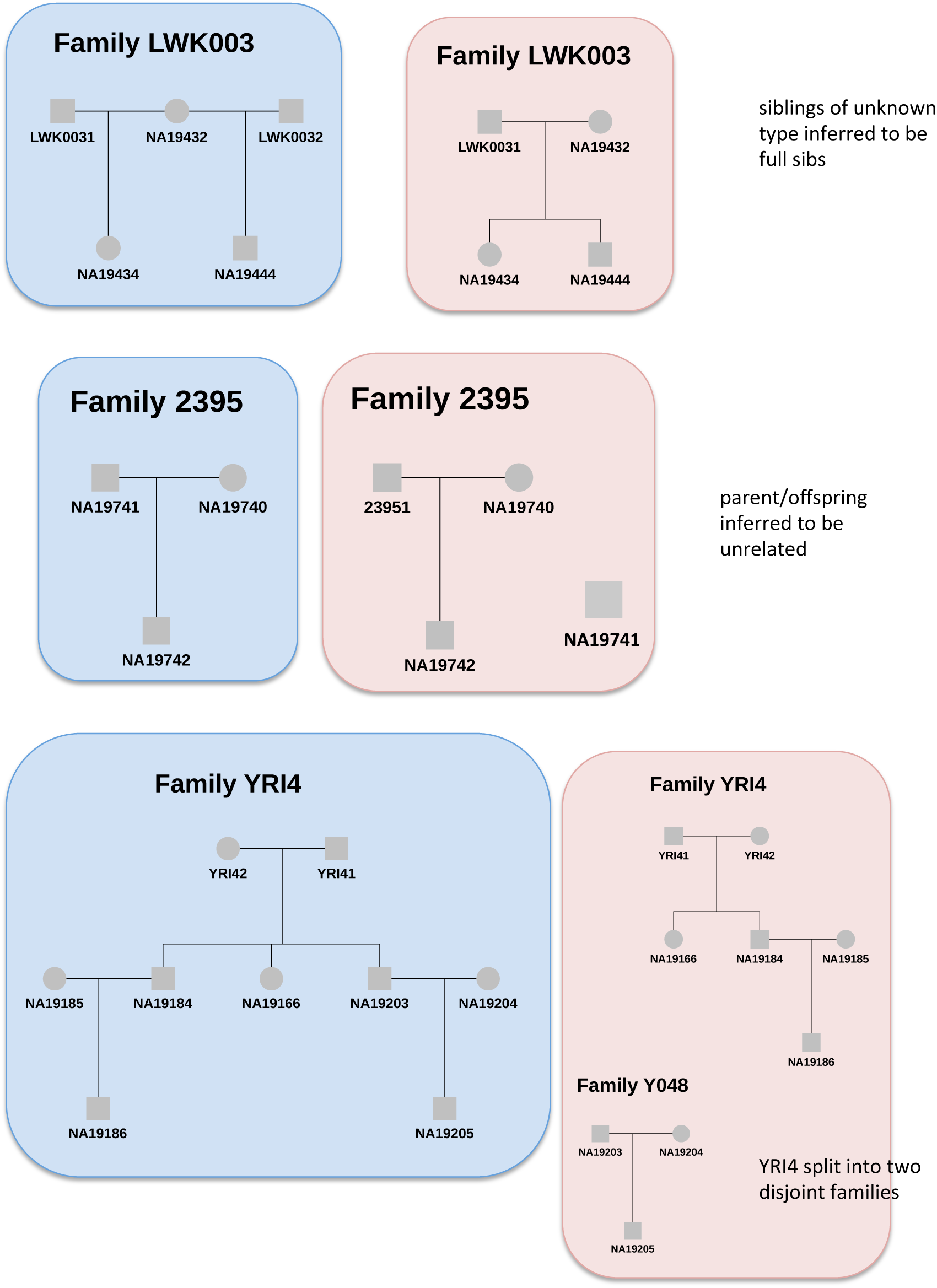

